# Bioefficacy of hydroxytyrosol-rich food supplements on preventing lipid peroxidation in healthy men

**DOI:** 10.1101/2022.09.21.508834

**Authors:** Cecilia Bender, Ilaria Candi, Eva Rogel

**Affiliations:** Institut Kurz GmbH; Stöckheimer Weg 1, 50829 Köln, Germany; Istituto Kurz Italia S.R.L; Via Golfo dei Poeti 1/A, 43126 Parma, Italy

**Keywords:** olive phenolics, hydroxytyrosol, oxidized LDL, ox-LDL, isoprostanes, F2-isoprostanes

## Abstract

In the present study we report the efficacy of food matrices derived from olives in preventing oxidation of low-density lipoprotein cholesterol (LDL) and lipid peroxidation. To this end, 12 healthy volunteers were divided into 3 groups and cross-received a single dose of olive phenolics, mainly hydroxytyrosol (HT), provided as a liquid dietary supplement (30.6 or 61.5 mg HT), or as fortified extra virgin olive oil (12.19 mg HT + tyrosol). Blood and urine samples were collected at baseline and up to 12 hours after ingestion. Plasma oxidized LDL levels were measured by ELISA using a monoclonal antibody, while F2-isoprostanes were quantified in urine by UHPLC-MS/MS. Despite the great variability between individuals, a tendency to reduce lipoxidation reactions has been observed after olive phenolics intake in both, blood and urine. In addition, the subgroup of individuals with the highest baseline lipoxidation level showed a decrease in F2-isoprostanes (p < 0.05) after taking the food supplements, as well as a marked decrease in oxidized LDL levels (p < 0.01) after intake of the food supplement with the lowest HT dose. These promising results suggest that HT supplementation could be a useful aid in preventing lipoxidation. Additionally, people with a redox imbalance could benefit even more from supplementing with bioavailable HT.

## 1. Introduction

An altered balance of free radicals, resulting in high levels of oxidative stress, is associated with many chronic human diseases such as cardiovascular diseases, a leading cause of global mortality, including atherosclerosis [1, 2, 3]. The first biochemical event in atherosclerosis is the oxidation of LDL in the vascular wall producing oxidized-LDL (oxLDL) [4], the latter involved in the formation and progression of atherosclerotic plaque [5-6]. The oxidative process of the LDL consists in a chemical modification of the protein moiety through the myeloperoxidase-derived enzyme [7], or the oxidation of LDL’s polyunsaturated fatty acids (PUFA) [8]. Nutrition plays an important role in the prevention of atherosclerosis [9], in particular dietary antioxidants can protect LDL from oxidation [10-11].

The olive tree contains phenolic compounds, such as HT, tyrosol (Tyr) and oleuropein (Ole) [12-13], which have key roles in plant physiology such as enhancing resistance to insects and microorganisms [14]. The oil obtained from the pressing of olive fruits contains phenolic compounds that contribute to the protection of LDL from oxidation. Different qualities of olive oils are available on the market from which only extra virgin olive oil (EVOO), the lesser processed olive oil which is obtained from milling and cold pressing, may contains enough phenolics [15-16] such as to protect LDL from oxidation.

In particular the HT, one of the main antioxidant in olives, is also believed to act in table olives and oil, and has received health claim approval in the EU [17]. Indeed, the European Food Safety Authority (EFSA) has confirmed that 20 grams of olive oil containing at least 5 mg of HT and its derivatives (Tyr and Ole complex) contribute to the protection of LDL from oxidation.

Despite the belief that a diet rich in olive oil would guarantee an adequate intake of olive phenolics, which due to their antioxidant capacity, would give a positive effect in the prevention of cardiovascular risks by preventing the LDL oxidation, other evidence suggests that the daily intake of HT, Tyr and phenolic compounds would be insufficient [18-19] as to obtain the desired physiological effect. Since these compounds are present in the olive fruit, they are also found in olive derivatives such as olive oil and vegetation waters. And since many of these substances are hydrophilic, they are found to a greater extent in the aqueous fraction that results from the pressing of the olive fruit during oil production [20]. It seems so that to ensure the minimum useful daily dose for the protection of LDL, it would be necessary to enrich the olive oil with HT, alternatively, the consumption of food supplements containing bioavailable HT would be a valid resource.

In this context, we have recently reported cellular antioxidant properties of two dietary supplements rich in natural HT derived from olives [20], one with the addition of 6% lemon juice and the other with 70% grape juice. The results derived from this preliminary study denoted an important antioxidant potential of both supplements in cellular models. More recently, in a controlled clinical study with 12 healthy men, we have demonstrated the bioavailability of HT contained in the same aqueous food supplements [21]. As a secondary outcome of this clinical study, we report here the effect of both food supplements in reducing oxidative damage by measuring two reliable markers of lipoxidation, the oxLDL in blood and the F2-isoprostanes (F2-IsoPs) in urine.

F2-IsoPs are prostraglandin-like lipid peroxidation products of the arachidonic acid, the main PUFA present in human cells. Numerous studies showed these molecules to be accurate markers for systemic oxidative damage in plasma or urine [22-24] and their decrease indicate a lower in vivo peroxidation of lipids.

Our data indicate that a single intake of HT through the dietary supplements evaluated tends to reduce the mean concentration of oxLDL in blood. In subjects whose oxLDL concentrations were particularly elevated at baseline, the decrease after ingestion was statistically significant. In addition, the F2-IsoPs measurement from these latter subjects show that the intake of either food supplement significantly reduces the F2-IsoPs content in urine.

Despite the small sample size, the results obtained here are promising, especially for the balance of redox status in the body. Quite interesting is the marked decrease of both biomarkers of oxidative stress as a result of taking these HT-rich food supplements, with the caveat that the effect depends on the individual’s baseline condition, people who have high starting levels of these markers will experience higher benefit.

## 2. Materials and Methods

### Standards and Reagents

Citric acid, phosphoric acid, L(+)-ascorbic acid, 1-butanol and ethylacetate were from Roth (Karlsruhe, Germany). Oxidized LDL ELISA kits (Cod: 10-1143-01) were from Mercodia (Uppsala, Sweden). 9α,11α,15S-trihydroxy-prosta-5Z,13E-dien-1-oic-3,3,4,4-d4 acid (PGF2α-d4, CAS 34210-11-2), 9β,11β,15R-trihydroxy-(8β,12α)-prosta-5Z,13E-dien-1-oic acid (ent-PGF2α, CAS 54483-31-7), 9α,11α,15S-trihydroxy-(8β)-prosta-5Z,13E-dien-1-oic acid (8-iso PGF2α, CAS 27415-26-5), and (3Z)-5-[(1S,2R,3R,5S)-3,5-dihydroxy-2-[(1E,3S)-3-hydroxy-1-octen-1-yl]cyclopentyl]-3-pentenoic acid (2,3-dinor-8-isoPGF2α, CAS 221664-05-7) were purchased from Cayman Chemical (Ann Arbor, MI, USA). Formic acid was from Merck (Darmstadt, Germany). LC-MS-grade water and methanol were purchased from VWR Chemicals (Darmstadt, Germany). HTEssence® Hydroxytyrosol Liquid was kindly provided by Wacker Chemie AG (Munich, Germany).

### Investigational Products (IPs)

Liquid food supplements and the EVOO were provided by Fattoria La Vialla S.A.S (Castiglion Fibocchi, Arezzo, Italy). Food supplements are derived from olive fruit (*Olea europaea L*.) vegetation water subjected to filtration and concentration, commercial brands are Oliphenolia® bitter and Oliphenolia®, hereafter referred to as IP-1 and IP-2, respectively. IP-1 consists of a 94% concentrated olive aqueous fraction and 6% concentrated lemon juice (*Citrus limon L*. fructus), containing 30.6 mg HT and 0.04 mg Ole per portion. While IP-2 contained a higher dose of HT (61.5 mg HT: 0.07 mg Ole) and is characterized by 30% of further concentrated olive extract and 70% concentrated grape (*Vitis vinifera L*. fructus) juice. Quantification of HT and derivatives in the food supplements was conducted as previously described [20]. Additionally, EVOO was spiked with HT to reach a final concentration of 5.77 mg/20 gr (equivalent to 12.19 mg of HT + Tyr), and assigned as a comparison group. Final concentration in the fortified EVOO after acid hydrolysis was conducted via HPLC-UV by ADSI (Austria).

### Participants and study design

The study protocol was approved by the Ethics Committee of State Medical Association of Rheinland-Pfalz (Mainz, Germany). The clinical study was conducted as set out in the Code of Ethics of the World Medical Association (Declaration of Helsinki) by daacro GmbH & CO at the Science Park Trier (Germany) and is registered at ClinicalTrials.gov (identifier: NCT04876261). Written informed consent was obtained from all participants prior to starting the trial. Twelve healthy male volunteers were recruited who met the inclusion and exclusion criteria (age 21-50, BMI> 18.5 <29.9 kg / m^2^, no-smokers, no-eating disorder, no-drug treatment in the previous or ongoing 2 weeks, no intake of food supplements, no drug or alcohol abuse). Diet indications included avoid consuming olive derived products as well as alcohol and supplements with HT, vitamins, minerals and antioxidants, 2-4 days before the first intake and during the whole study. In addition, three days prior to and at each intervention, volunteers avoided moderate or intense physical activity.

Study design was single-blind, randomized, single-dose, three way cross-over, in which volunteers ingested different concentrations of olive phenolics through a single dose of the corresponding IPs. IPs were orally administrated together with 200 mL of water after an overnight fast of at least 10 hours (hrs). Volunteers underwent a wash-out period of 6 days between the interventions to avoid interference between the IPs.

### Sampling

At each intervention visit a baseline blood sample was collected immediately before the administration of the IP. Further blood samples were collected 0.5, 1, 1.5, 2, 4 and 12 hrs after the intake. EDTA-plasma samples were obtained and stored at -80 °C until analysis.

At each intervention visit a baseline urine sample was collected from -240 to 0 min before the administration of the IP. Further urine samples were collected after the intervention from 0 to 30 min, 30 min to 1 hr, 1 to 1.5 hr, 1.5 to 2 hrs, 2 to 4 hrs, and 4 to 12 hrs. Total volumes excreted were measured, stabilized with 1.88 g/L of ascorbic acid, and stored at −80 °C until analysis.

### Analysis of plasma oxLDL

Plasma samples were thawed at room temperature and immediately quantified for oxLDL by a sandwich ELISA assay according to the manufacturer’s recommendations. Briefly, the sandwich assay uses two monoclonal antibodies against separate antigenic determinants of the oxidized apolipoprotein B molecule. During a first incubation the plasma oxLDL reacts with the capture antibody mAb-4E6. The next steps include incubation with the peroxidase-conjugated secondary anti-human apolipoprotein B antibody, which is detected spectrophotometrically by reaction with 3,3 ‘, 5,5’-tetramethylbenzidine. oxLDL concentration was recorded in duplicates in a Fluostar OPTIMA reader (BMG Labtech, Offenburg, Germany) and calculated with a Five Parameter Logistic (5PL) curve and automatic weighting using 1 / Y^2^. The mean values of oxLDL, expressed in units per liter (U/L), were used for the statistical analysis.

### Analysis of urinary F2-IsoPs

Isomers 8-isoPGF2α, ent-PGF2α, and 2,3-dinor-8-isoPGF2α were evaluated by UHPLC-MS/MS following an internal method based on unpublished work. In brief, urine samples were thawed and spiked with formic acid and of PGF2α-d4. Extraction solvent (5% butanol in ethylacetate) were added, mixed and placed in ice bath for 3 min, following by centrifugation. The organic phase was collected and evaporated under nitrogen flow. Extraction step was repeated twice. Dry sample was resuspended with methanol and formic acid, filtered through 0.2 μm regenerated cellulose filters (Macherey Nagel, Düren, Germany) and transferred to a new vial.

Measurement was conducted with an Acquity UPLC I-Class system coupled to a XEVO-TQS micro mass spectrometer (Waters, Milford, MA, USA) using standard substances as reference. The instrument consisted of a sample manager cooled at 10 °C, a binary pump, a column oven, and a diode array detector. The column oven temperature was set at 40 °C. Eluent A was acetonitrile with 0.1% formic acid, eluent B was water with 0.1% formic acid. The flow was 0.4 mL/min on a BEH Shield RP18 column (Acquity, 150 mm x2.1 mm, 1.7 μm particle size) combined with a BEH Shield RP18 precolumn (Acquity, 2.1 mm × 5 mm, 1.7 μm), both from Waters (Milford, MA, USA). The gradient started with 30% A and raised to 80 %. The peaks were identified by MS/MS.

All samples were run in duplicate. Data were acquired and processed using MassLynx (Waters, Milford, MA, USA) and normalized by dividing the concentration by the urinary creatinin content in the sample. Mean values of F2-isoP isomers, expressed in μg/gr, were used for the statistical analysis.

### Data analysis

The average values of concentration for each of the samples were calculated in Microsoft Excel version 16.0. Average values were further processed using GraphPad version 5.00 (San Diego, California, USA) software to represent in graphs and table. For the raw statistics, classical statistical methods using median, mean, standard deviation and confidence intervals were used. Data are expressed as mean ± standard error (SEM) otherwise indicated. Student’s paired t-test was performed to compare the results before and after intake of each IPs, and to compare the results before and after the ingestion of HT irrespective of the product.

## 3. Results

### 3.1 Plasma oxLDL

OxLDL was quantifiable in all plasma samples collected before (0 hr) and after HT intake through IP-1, IP-2, and EVOO. Figure 1 shows box plots representing the oxLDL content before HT ingestion and up to 12 hrs after ingestion. A significant reduction in oxLDL was observed after 1 hr (p<0.05) and 2 hrs (p<0.01) of intake, regardless of food matrix (Figure 1A). Furthermore, when the statistical analysis is stratified to include only the interventions with the dietary supplements, the oxLDL is significantly lower (p<0.05) from baseline after 1 hr of the intake (Figure 1B).

**Figure 1:**
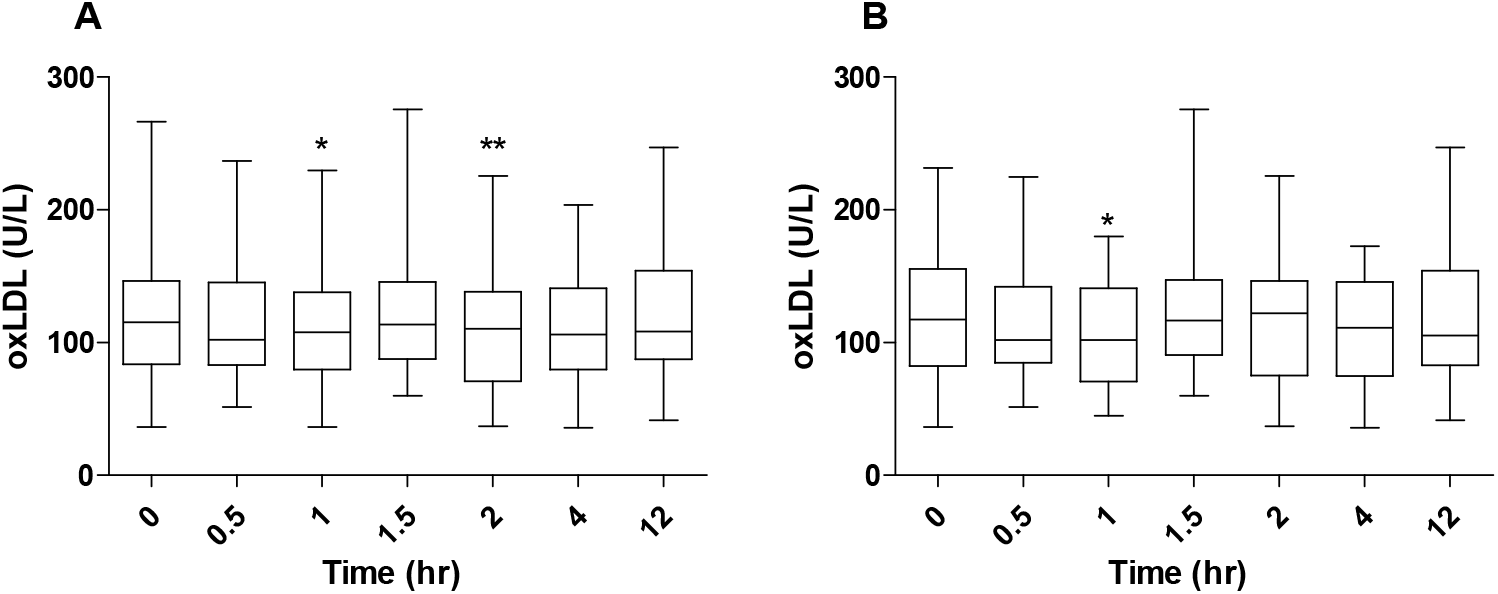
oxLDL values before and after treatment with HT. The graph shows the median oxLDL concentration (U/L) in plasma, maximum and minimum values, 25% and 75% percentiles. **(A)** oxLDL values over time regardless of the HT source. **(B)** oxLDL values before and after HT administration through food supplements. *: significantly different from baseline with a p<0.05, **: significantly different from baseline with a p<0.01.

The analysis of the average oxLDL of the independent interventions shows a trend towards the reduction of oxLDL after supplying HT (Figure 2), however the small sample size and the wide inter-individual variation observed do not allow to show statistical significance. In particular, the intervention with IP-2, containing the higher dose of olive phenolics (61.5 mg HT: 0.07 mg Ole), slowly decreased the mean oxLDL, reaching the lowest point one hr after its ingestion and values similar to the initial ones at 4 and 12 hrs post-treatment (Figure 2A). IP-1, containing 30.6 mg HT: 0.04 mg Ole, shows an apparent two-step decrease in mean oxLDL (Figure 2B), respectively at 1 and 4 hrs post-intervention. While the administration of EVOO, containing 12.19 mg HT and Tyr, shows a slow decrease in mean oxLDL (Figure 2C), reaching the lowest point 2 hrs after intake and then increasing until reaching the maximum value 12 hrs after the intervention.

**Figure 2:**
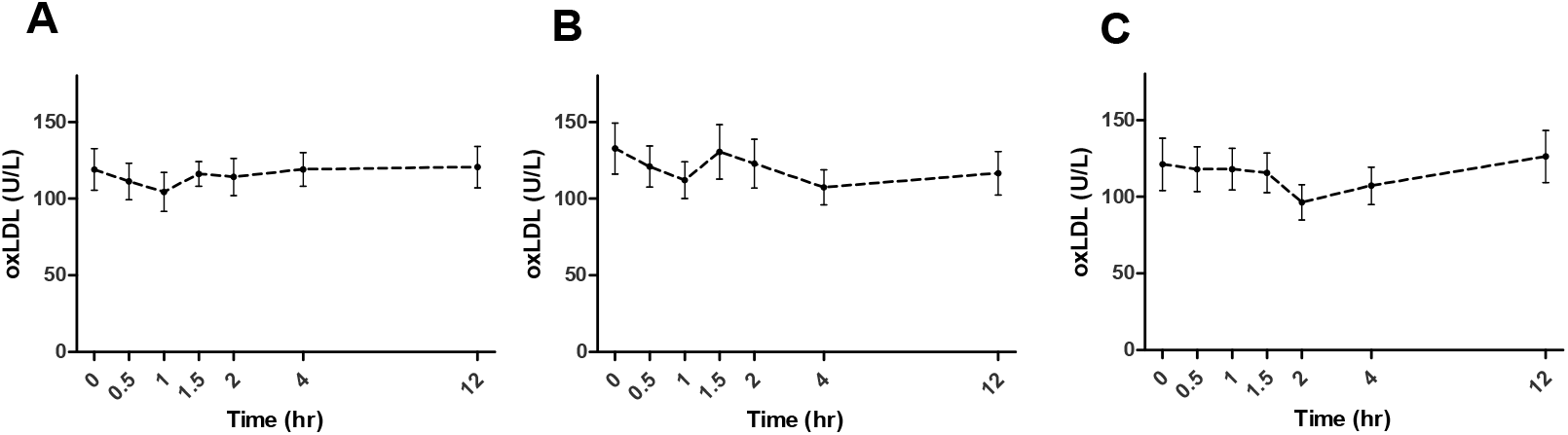
Mean plasma oxLDL concentration over time before (0 hr) and after dietary intervention. **(A)** IP-2, **(B)** IP-1, **(C)** EVOO. Bars: SEM

It is interesting to note that, by applying an oxLDL cut off of 101 U/L before the interventions, that is mean values higher than reported for healthy subjects [25], the oxLDL was significantly reduced after 1, 2, 4, and 12 hrs of IP-1 intake, and after 2 hrs of EVOO intake (Figure 3). This observation suggests that the HT protection of LDL against oxidation is likely to be more effective in individuals with an imbalance in the redox status.

**Figure 3:**
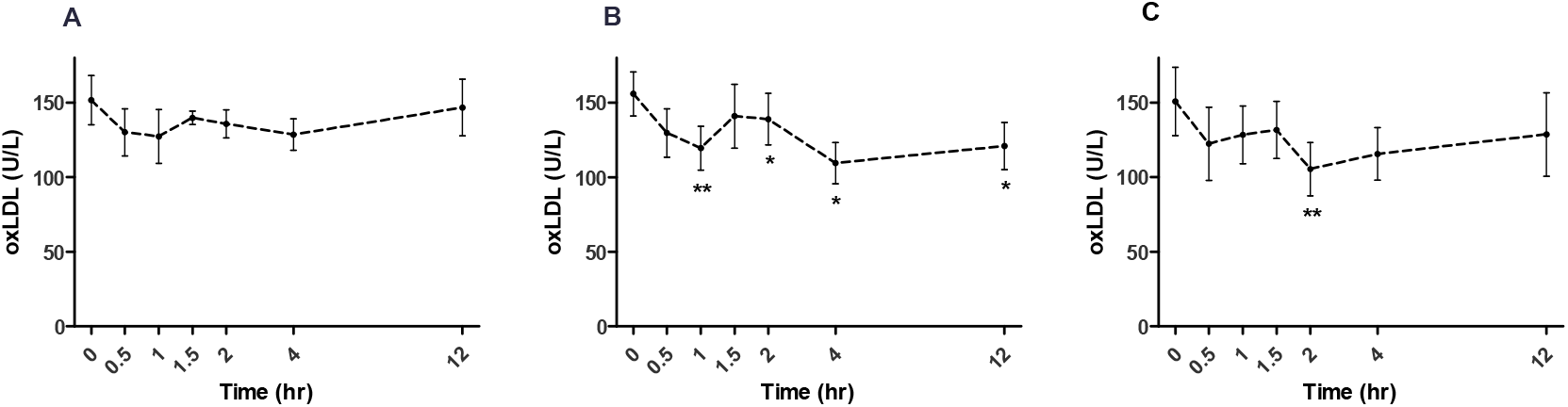
Mean oxLDL plasma values over time before (0 hr) and after ingestion of **(A)** IP-2, **(B)** IP-1, and **(C)** EVOO using the highest oxLDL concentration at baseline (cut off 101 U/L). Bars: SEM. *: p<0.05, and **: p<0.01 versus baseline.

### 3.2 Urinary F2-IsoPs

In addition to the plasma oxLDL, the degree of oxidative stress was evaluated in urine by quantification of F2-IsoPs. Three different isomers, namely 8-isoPGF2α, ent-PGF2α, and 2,3-dinor-8-isoPGF2α were quantified at baseline in all the participants, while after the intake of the HT-rich food supplements they were quantified only in those subjects showing high lipoxidation levels at baseline. After the intake of either of the two food supplements it was shown a significant decrease on F2-IsoPs (Table 1). Quantitatively, the excreted amount of F2-IsoPs (as the sum of 8-isoPGF2α, ent-PGF2α, and 2,3-dinor-8-isoPGF2α isomers) resulted higher at baseline compared with after the intake of the food supplements, being more pronounced for IP-2 compared with IP-1.

**Table 1:**
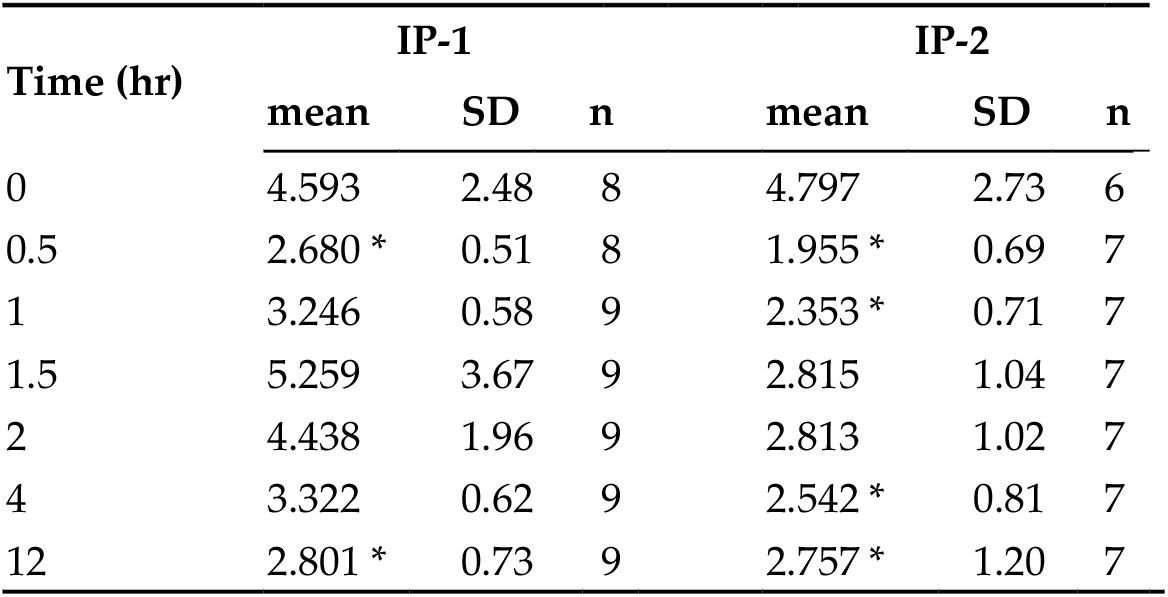
F2-IsoPs excretion following HT-rich food supplements intake in subjects with high level at baseline. Mean content (μg/gr) is expressed as the sum of 8-isoPGF2α, ent-PGF2α and 2,3-dinor-8-isoPGF2α. SD: standard deviation; n: sample size; *: p<0.05 versus baseline.

| Time (hr) | IP-1 |  |  | IP-2 |  |  |
| --- | --- | --- | --- | --- | --- | --- |
|  | mean | SD | n | mean | SD | n |
| 0 | 4.593 | 2.48 | 8 | 4.797 | 2.73 | 6 |
| 0.5 | 2.680 * | 0.51 | 8 | 1.955 * | 0.69 | 7 |
| 1 | 3.246 | 0.58 | 9 | 2.353 * | 0.71 | 7 |
| 1.5 | 5.259 | 3.67 | 9 | 2.815 | 1.04 | 7 |
| 2 | 4.438 | 1.96 | 9 | 2.813 | 1.02 | 7 |
| 4 | 3.322 | 0.62 | 9 | 2.542 * | 0.81 | 7 |
| 12 | 2.801 * | 0.73 | 9 | 2.757 * | 1.20 | 7 |

In particular, changes in the isomers 8-isoPGF2α (Figure 4B and 4E) and ent-PGF2α (Figure 4A and 4D) were significantly lower after the administration of both nutritional supplements (p<0.05) and were more pronounced for IP-2, the food supplement with the higher HT-dose. While 2,3-dinor-8-isoPGF2α changes resulted significantly lower (p<0.01) soon after the intake of IP-2 (Figure 4C) but not significant after the intake of IP-1.

**Figure 4:**
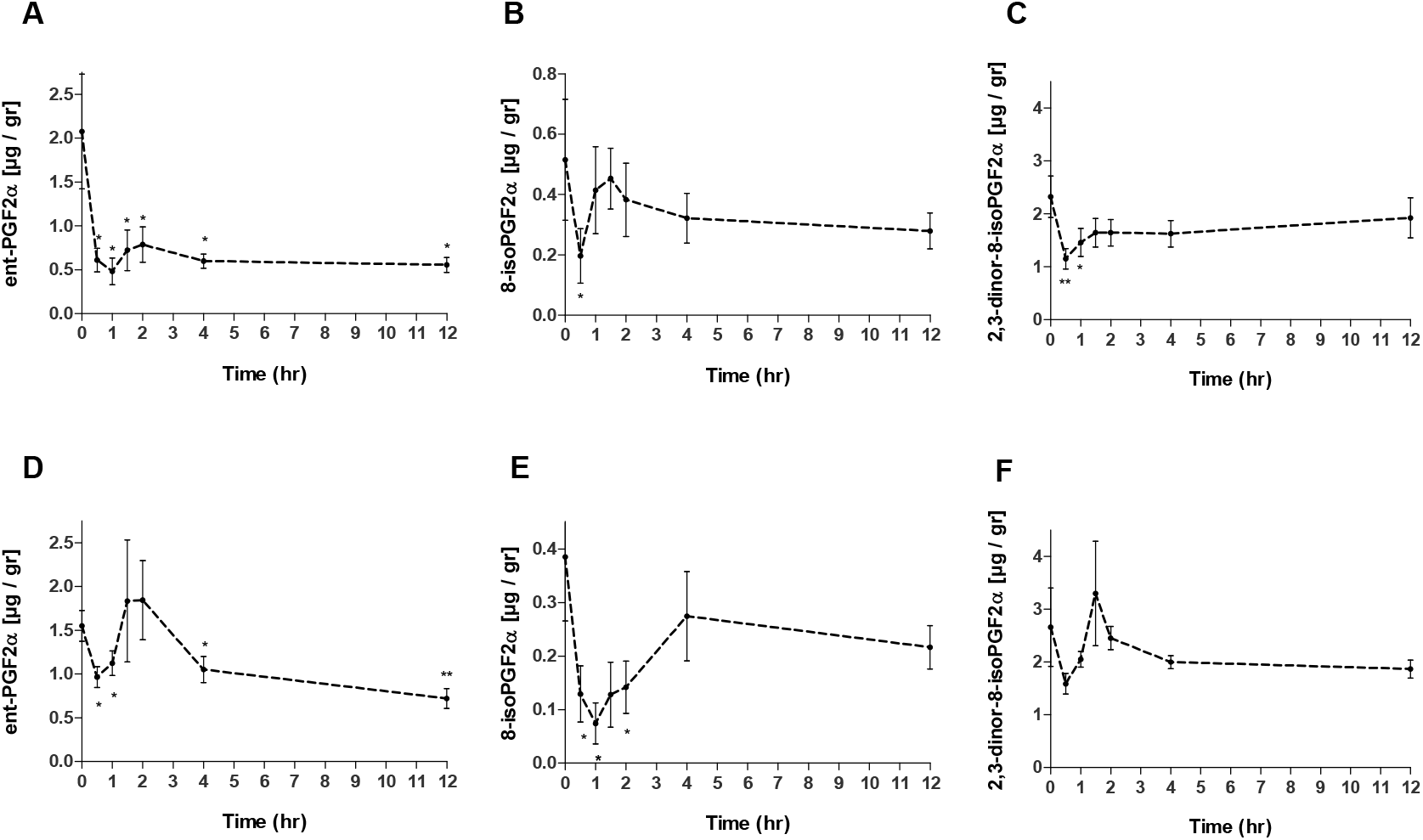
Mean urinary IsoPs content over time, before (0 hr) and after ingestion of IP-2 **(A, B, C)** and IP-1 **(D, E, F)** using the highest IsoPs levels at baseline. Bars: SEM. *: p<0.05, and **: p<0.01 versus baseline.

## 4. Discussion

The objective of this crossover study was to verify whether two aqueous food supplements, rich in olive phenolic compounds, particularly HT, are effective in preventing lipid peroxidation in humans. To this aim, plasma oxLDL and urinary F2-IsoPs were used as reliable biomarkers.

It has been well documented that olive oil phenolics reduce oxLDL in vivo after short-medium term interventions [26-34]; however to the best of our knowledge there are no reports showing that also olive phenolics provided in a watery matrix can reduce oxLDL in vivo. Our results show a significant reduction in plasma oxLDL as soon as after 1 and 2 hrs of intake of a single dose of olive phenolics regardless the food matrix. We also observed a downward trend (p>0.05) in the level of oxLDL in each of the three groups studied separately, and a significant reduction at 1 h of the intake of aqueous food supplements.

Furthermore, considering only subjects with high levels of oxLDL at baseline, a significant reduction in oxLDL was observed after the administration of IP-1 (30.6 mg HT + 0.04 mg Ole) and EVOO (12.19 mg HT + Tyr), respectively at 1, 2, 4 and 12 hrs for IP-1 and after 2 hrs for the fortified EVOO. This finding reinforces the notion that people with low oxidation levels rarely see much benefit from supplementing antioxidative stress nutrients. Furthermore, this result suggests that the oxLDL-reducing effect is even more effective using a medium water-based concentration than a low concentration in fortified oil.

It is worth mentioning that other compounds present in the food supplements may interfere with LDL-protection efficacy, as both supplements contain concentrated aqueous olive extract at different doses along with grape concentrate (IP-2: 61.5 mg HT + 0.07 mg Ole) or lemon juice (IP-1: 30.6 mg HT + 0.04 mg Ole). Unlike what was reported in the literature [27-31] for olive oil intake, our results do not show a simple dose-dependence of an oxLDL protective effect. In fact, IP-2, the food supplement containing the highest dose of olive phenolics, did not cause a higher oxLDL reduction than IP-1, which contains almost half the HT-dose together with the lemon juice concentrate. Neither was IP-2 more efficient than the fortified EVOO.

The results obtained considering only subjects with a high oxLDL level at baseline suggest that the olive phenolics efficacy is particularly effective on individuals who are experiencing an oxidative imbalance. This observation is supported by several human trials conducted in subjects with high oxidative stress condition, in which a significant reduction of lipoxidation was documented by using different biomarkers, such as plasma oxLDL or urinary IsoPs [34-38]. In particular, Sarapis [34] recently reported a significant decrease of oxLDL consuming high dose of olive oil polyphenols, reduction that became more pronounced in subjects with high cardiometabolic risk.

In 2011, the European Food Safety Authority (EFSA) published a scientific opinion in which it is stated that a daily consumption of at least 5 mg of HT and derivatives (Tyr and Ole) contained in olive oil will protect the LDL against oxidation [17]. Nevertheless, other researchers pointed out that the daily intake of HT, Tyr and phenolic compounds through a typical mediterranean diet would supply only around 2 mg [18]. Moreover, the intake of Tyr and HT from virgin olive oil would be between 88.5 and 237.4 μg daily [19]. These studies seem to indicate that the amount of HT and its derivatives ingested daily with the mediterranean diet or with olive oil alone would be insufficient to reach the EFSA-stated minimum intake-level of 5 mg. Therefore, it seems to be advisable to increase the intake of HT and its derivatives in order to obtain the protective effect of LDL. Such increase of intake can be achieved by supplementation with HT-enriched oil. Alternatively it can be even better achieved by consumption of HT-containing supplements. Although EFSA’s statement refers to the HT consumed in 20 g of olive oil, in this study we show that both food supplements on watery basis containing bioavailable HT cause a higher effect as compared to EVOO.

In addition to plasma LDL oxidation, another important marker to consider for the prevention of oxidative processes in vivo is the F2-IsoPS. Few studies conducted with healthy volunteers have evaluated the IsoPs content after ingesting foods derived from olives. The results remain somewhat contradictory.

On the one hand, a reduction in urinary 8-iso-PGF2α was observed inversely proportional to the phenolic content of olive oils (phenolic concentrations between 24.38 and 97.5 mg / dose) when administered in a single dose [39].

On the other hand, further short-term studies found no significant changes in F2-IsoPs excretion. In a crossover study with olive oil rich in phenolics conducted with 182-184 healthy volunteers, in which despite the higher dose of phenolics tested (366 mg/kg, 25 mL daily, 21 days) decreased the oxLDL, no such an effect was observed on IsoPs in plasma [30]. However, a significant decrease in plasma IsoPs was observed for the above study when comparing only the baseline data and the endpoint data at the end of the crossover interventions [40]. Regarding the studies with dietary supplements, no changes in urinary levels of F2-IsoPs have been recorded in young people after multiple dosages of olive leaf supplements in liquid or capsule formats [41].

Although human trials on the F2-IsoPs modulation from olive-derived foods have yielded contradictory results, it is noted that the trials described above evaluated different doses, population, duration, and food matrix, as well as different analytical methods and biological fluids were analyzed, all variables that affect the final outcome [42]. Furthermore, it is recognized that even under normal conditions IsoPs appear in the plasma and urine and their levels are only amplified by oxidative stress [43-44]. For this reason they are used as markers of lipid peroxidation in both human and animal models. Indeed, several clinical studies that report a significant modulation of IsoPs levels after a specific treatment, have been carried out in subjects showing oxidative stress condition [36, 38, 45-52]; that is, the reduction of oxidative markers could be significant only if their levels are high at the beginning of the study. In the contrast to this, antioxidant supplementation in subjects with a balanced redox state seems to be of little clinical relevance.

In the present study we evaluated the degree of urinary excretion of F2-IsoPs in healthy participants before treatments, and after taking the HT-rich food supplements only in those subjects with a high level of oxidation at time zero. Applying this criterion, we found that both dietary supplements significantly reduced the F2-IsoPs excreted in the urine, thus suggesting that subjects with redox imbalance may benefit of the food supplements integration to protect lipids from peroxidation.

## 5. Conclusions

The present study shows that the antioxidant effects of food supplements rich in olive phenolic compounds and, in particular, in bioavailable HT, are promising for the reduction of lipooxidation in vivo. An oxLDL reduction tendency was observed shortly after intake of the IPs. Moreover, oxLDL was significantly reduced in plasma after 1 and 2 hrs of olive phenolic compounds intake, regardless of the food source that contained it (aqueous or oily), and also significantly reduced after 1 hr of the intake of the watery food supplements. In general, these results support the positive effect of olive phenols in the prevention of lipid peroxidation. We further support the finding that antioxidant effects from antioxidative food can be expected most from people with elevated oxidative stress. Under this condition the performance of IP-1 (30.6 mg HT : 0.04 mg Ole) was superior than that of the EVOO (12.19 mg HT + Tyr) and also higher than IP-2.

By measuring the F2-IsoPs in urine we showed that in healthy subjects with high oxLDL values at the beginning of the study, both food supplements significantly reduced the excretion of F2-IsoPs.

Overall, the results obtained in this study indicate the efficacy of both food supplements to reduce the level of lipid peroxidation soon after intake, which is a valuable aid to prevent and combat oxidative damage in the human body.

## Author Contributions

Conceptualization, methodology, investigation, and data curation, C.B.; formal analysis, C.B., I.C., and E.R.; original draft C.B., and I.C.; writing—review and editing, C.B.; visualization, E.R.; All authors have read and agreed to the published version of the manuscript.

## Institutional Review Board Statement

The study was conducted in accordance with the Declaration of Helsinki, approved by the Ethics Committee of State Medical Association of Rheinland-Pfalz (Mainz, Germany), and registered at ClinicalTrials.gov (identifier: NCT04876261).

## Informed Consent Statement

Informed consent was obtained from all subjects involved in the study.

## Data Availability Statement

Not applicable.

## Acknowledgments

The authors would like to acknowledge Dr. Christian Golz and Dr. Helmut H. Weidlich for their support in interpreting the data, Dr. Sarah Strassmann for technical support in validating the IsoPs method, and Mrs. Übelhör on whose unpublished master thesis the method is based.

